## Supplemental figures and legends for "The cation channel mechanisms of subthreshold inward depolarizing currents in the VTA dopaminergic neurons and their roles in the chronic-stress-induced depression-like behavior"

**sFig. 1** The targeted regions for VTA DA projection.

**A** and **B**. Injection sites (red colored) in sagittal (A) and coronal plane (B) of mouse brain with DAPI (405 nm)-counterstained sections (80 μm) showing locations of retrobeads (546 nm, yellow). From top to bottom: mPFC (bregma +2.22 mm), BLA (bregma -1.58 mm)，NAc ms (bregma +1.10 mm),  NAc c (bregma +0.98 mm), NAc ls (bregma +0.86 mm).

**sFig. 2** The subregions of projection-specific VTA DA neurons.

**A.** Confocal images showing the anatomical distribution of retrobeads (red) in the VTA after DAT-immunofluorescence (green) staining at different magnification (i, 20×; ii, 40×); individual labeled cells are shown in (iii). Scale bars i = 100 μm, ii = 20 μm and iii = 10 μm. **B** and **C.** Average retrogradely labeled cell distributions in the VTA.

**sFig. 3** Blocking HCN channels does not affect the spontaneous firing of the VTA DA neurons in adult male mice.

**A** and **B.** Effects of HCN channel inhibitors CsCl (A) and ZD7288 (B) on the sag membrane potential of adult male VTA DA neurons. The sag membrane potential was recorded using whole-cell current-clamp method, and sag was calculated as the difference between the peak negative voltage and steady-state negative voltage responding to hyperpolarizing current injection (-100 pA). Ai and Bi: CsCl (3 mM, n = 4, N = 4) and ZD7288 (60 μM, n = 4, N = 4) significantly decreased the sag potentials. Aii and Bii: Summary for CsCl and ZD7288 induced inhibition on sag potentials. Paired-sample T test, CsCl: t = 4.246, df = 3, 95% CI: -22.03 to -3.155, *$P$ = 0.0239; ZD7288: t = 5.017, df = 3, 95% CI: -24.83 to -5.555, *$P$ = 0.0152. **C** and **D.** Effects of HCN channel inhibitors CsCl (C, n = 7, N = 4) and ZD7288 (D, n = 13, N = 5) on the spontaneous firing activity of the VTA DA neurons (DAT positive). The spontaneous firing of the VTA DA neurons was recorded using loose cell-attached current clamp method on the brain slice of the VTA. The example time-courses (i) and the summarized data (ii) for the effects of CsCl and ZD7288 were shown. Paired-sample T test, CsCl: t = 0.7171, df = 6, 95% CI: -0.4325 to 0.7911, $P$ = 0.5003; ZD7288: t = 1.349, df = 12, 95% CI: -0.4033 to 0.09486, $P$ = 0.2022, n.s. $P >$ 0.05, *$P <$ 0.05. n is number of DAT-positive neurons recorded and N is number of mice

**sFig. 4** HCN channel blocker does not affect the spontaneous firing of the adult male VTA DA neurons projecting to NAc lateral shell but block the spontaneous firing of the VTA DA neurons bathed in a low extracellular $K^+$ which hyperpolarized the resting membrane potential.

**A** and **B.** Effects of HCN channel inhibitors (CsCl A, ZD7288 B) on the spontaneous firing activity of the VTA DA neurons projecting to NAc lateral shell. The spontaneous firing of VTA DA neurons were recorded using loose cell-attached current clamp method on the brain slice of the VTA. The example time-courses (i), traces (ii) and the summarized data (iii) for the effects of CsCl (A, n = 5, N = 4, Red and DAT positive), ZD7288 (B, n = 5, N = 5, Red and DAT positive) were shown. Paired-sample T test, CsCl: t = 0.4798, df = 4, 95% CI: -0.7601 to 0.5361, *P* = 0.6564; ZD7288: t = 0.4421, df = 4, 95% CI: -0.4368 to 0.3168, *P* = 0.6813. iv: a map of a coronal midbrain slice indicating the location of red neurons that were recorded and subsequently identified as DA neurons which were DAT positive with single cell-PCR (red dots). **C.** i: Example time course for hyperpolarization of the resting membrane potential (RMP) induced by low extracellular $K^+$ (low $[K^+]_e$). ii: Summarized data for experiments shown in Ci (n = 7, N = 7), Paired-sample T test, t = 6.373, df = 6, 95% CI: -18.69 to -8.318, ****P* = 0.0007. iii: a map of a coronal midbrain slice indicating the location of neurons that were recorded and subsequently identified as DA neurons which were DAT positive with single cell-PCR (red dots). **D.** ZD7288 inhibited the spontaneous firing frequency of VTA DA neurons only in low $[K^+]_e$ condition. Example time courses of firing rates (i), traces (ii) and summarized data (iii) were shown (n = 7, N = 7, DAT positive). Wilcoxon matched-pairs signed rank test, W = -28.00, **P* = 0.0156. **E.** Example time-courses (i), traces (ii) and the summarized data (iii) for the effects of ZD7288 on the spontaneous firing of juvenile (less than 15 days postnatal) male VTA DA neurons were shown (n = 9, N = 5, DAT positive). Paired-sample T test, t = 3.345, df = 8, 95% CI: -1.487 to -0.2734, **P* = 0.0101, n.s. *P* > 0.05, **P* < 0.05, ****P* < 0.001. n is number of neurons recorded and N is number of mice.

**sFig. 5** Effects of TRPV channel blocker ruthenium red (RR) on the spontaneous firing of the male VTA DA neurons.

**A.** The inset on the top of i shows a map of a coronal midbrain slice indicating the location of neurons that were recorded and subsequently identified as DA neurons which were DAT positive with single cell-PCR (red dots). The bottom panel of i summarizes (n = 9, N = 6) the percentage of RR-activated and -inhibited VTA DA cells. ii: the summarized data for the effects of RR on the spontaneous firing frequency of VTA DA neurons. Paired-sample T test, t = 0.4486, df = 8, 95% CI: -0.7621 to 0.5139, $P = 0.6656$. **B** and **C.** Example time-course (i) and traces (ii) show the characteristic responses of firing activity to RR-activated (B) and -inhibited (C) neurons. n.s. $P > 0.05$, n is the number of DAT-positive neurons recorded and N is number of used mice.

**sFig. 6** The expression of TRPC4 and TRPC7 channels in male VTA TRPC6$^{+}$ DA neurons.

**A.** Single-cell PCR from 7 VTA TRPC6$^{+}$DAT$^{+}$ cells (C1-C7) and control. **B.** i and ii: Percentage of TRP channels (TRPC4 and TRPC7) positive neurons in the VTA TRPC6$^{+}$DAT$^{+}$ DA neurons.

**sFig. 7** Body weight and behavior of the male mice subjected to chronic mild unexpected stress (CMUS).

**A.** Changes in body weight of the CMUS mice from week 2 to 6 (Con: N = 10; CMUS: N = 10), Two-sample T test, t = 6.284, df = 18, 95% CI: -3.002 to -1.498, ****$P < 0.0001$, compared with the control mice. **B.** The total travel distance of the CMUS mice in the open-field test (Con: N = 10; CMUS: N = 10), Two-sample T test, t = 4.798, df = 18, 95% CI: -1767 to -690.9, ***$P = 0.0001$, compared with control mice. **C.** The time of the CMUS mice stayed in center zone in the open-field test (Con: N = 10; CMUS: N = 10), Two-sample T test, t = 4.733, df = 18, 95% CI: -27.29 to -10.51, ***$P = 0.0002$, compared with control mice. **D.** The percentage of time of CMUS mice stayed in open arm in the elevated plus-maze test (Con: N = 10; CMUS: N = 10), Two-sample T test, t = 4.440, df = 18, 95% CI: -12.36 to -4.421, ***$P$ = 0.0003, compared with control mice. **E.** The percentage of time of CMUS mice stayed in closed arm in the elevated plus-maze test (Con: N = 10; CMUS: N = 10), Mann-Whitney U test, U = 14, **$P = 0.0052$, compared with control mice. **F.** The immobility time of mice in the forced-swimming test (Con: N = 10; CMUS: N = 10), Two-sample T test, t = 4.570, df = 18, 95% CI: 17.56 to 47.44, ***$P$ = 0.0002, compared with control mice. **G.** The time of latency to fall in the rotarod test (Con: N = 10; CMUS: N = 10), Two-sample T test, t = 0.3323, df = 18, 95% CI: -11.18 to 15.38, $P$ = 0.7435, compared with control mice. n.s. $P$ > 0.05, **$P$ < 0.01, ***$P$ < 0.001, ****$P$ < 0.0001. N is the number of mice used.

**sFig. 8** Body weight and behaviors of the female mice subjected to chronic mild unexpected stress (CMUS).

**A.** The change in body weight of the CMUS female mice (Con: N = 10; CMUS: N = 10), Two-sample T test, t = 4.191, df = 18, 95% CI: -1.802 to -0.5984, ***$P$ = 0.0005, compared with control mice. **B.** Sucrose preference test for the CMUS female mice (Con: N = 10; CMUS: N = 10), Two-sample T test, t= 4.100, df = 18, 95% CI: -18.85 to -6.075, ***$P$ = 0.0007, compared with control mice. **C.** The total travel distance of the CMUS female mice in the open-field test (Con: N = 10; CMUS: N = 10), Two-sample T test, t = 5.552, df = 18, 95% CI: -1287 to -580.3, ****$P$ < 0.0001, compared with control mice. **D.** The time of the CMUS female mice stayed in center zone in the open-field test (Con: N = 10; CMUS: N = 10), Two-sample T test, t = 3.183, df = 18, 95% CI: -37.44 to -7.665, **$P$ = 0.0052, compared with control mice. **E.** The percentage of time of the CMUS female mice stayed in open arm in the elevated plus-maze test (Con: N = 10; CMUS: N = 10), Two-sample T test, t = 3.773, df = 18, 95% CI: -12.22 to -3.478, **$P$ = 0.0014, compared with control mice. **F.** The percentage of time of the CMUS female mice stayed in closed arm in the elevated plus-maze test (Con: N = 10; CMUS: N = 10), Two-sample T test, t = 2.966, df = 18, 95% CI: 3.143 to 18.40, **$P$ = 0.0083, compared with control mice. **G.** The time of latency to fall in the rotarod test (Con: N = 10; CMUS: N = 10), Two-sample T test, t = 0.7688, df = 18, 95% CI: -14.18 to 6.584, $P$ = 0.4520, compared with control mice. **H.** The immobility time of female mice in the forced-swimming test (Con: N = 10; CMUS: N = 10), Two-sample T test, t = 5.350, df = 18, 95% CI: 21.44 to 49.16, ****$P$ < 0.0001, compared with control mice. **I.** The immobility time of female mice in the tail suspension test (Con: N = 10; CMUS: N = 10), Two-sample T test, t = 5.308, df = 18, 95% CI:22.23 to 51.37, ****$P$ < 0.0001, compared with control mice. n.s. $P$ > 0.05, **$P$ < 0.01, ***$P$ < 0.001, ****$P$ < 0.0001. N is the number of mice used.

**sFig. 9** Expression of NALCN protein in the VTA of the CMUS male mice.

**A** and **B.** Western blot was used to measure the protein level of NALCN in the CMUS mice and control mice. Representative western blot assay showing the expression of NALCN and GAPDH in the VTA of the CMUS (N = 9) and control (Con, N = 12) mice (A), and the summarized data (B). Two-sample T test, t = 0.2724, df = 19, 95% CI: -0.2703 to 0.3512, $P = 0.7883$, n.s. $P > 0.05$. N is number of mice used.

**sFig. 10** Body weight and behaviors of the male mice subjected to Chronic restraint stress (CRS).

**A.** Experimental procedure timeline for CRS. **B.** The change in body weight of the CRS mice (Con: N = 10; CRS: N = 10), Two-sample T test, t = 5.481, df = 18, 95% CI: -2.144 to -0.9559, ****$P < 0.0001$, compared with control mice. **C.** Sucrose preference test for the CRS mice (Con: N = 10; CRS: N = 10), Two-sample T test, t = 4.653, df = 18, 95% CI: -19.05 to -7.199, ***$P = 0.0002$, compared with control mice. **D.** The total travel distance of the CRS mice in the open-field test (Con: N = 10; CRS: N = 10), Two-sample T test, t = 4.680, df = 18, 95% CI: -1385 to -526.6, ***$P = 0.0002$, compared with control mice. **E.** The time of the CRS mice stayed in center zone in the open-field test (Con: N = 10; CRS: N = 10), Mann-Whitney U test, U = 8, ***$P = 0.0007$, compared with control mice. **F.** The percentage of time of the CRS mice stayed in open arm in the elevated plus-maze test (Con: N = 10; CRS: N = 10), Mann-Whitney U test, U = 9, ***$P = 0.0010$, compared with control mice. **G.** The percentage of time of the CRS mice stayed in closed arm in the elevated plus-maze test (Con: N = 10; CRS: N = 10), Two-sample T test, t = 4.852, df = 18, 95%CI: 11.62 to 29.35, ***$P = 0.0001$, compared with control mice. **H.** The tail suspension test for the CRS mice (Con: N = 10; CRS: N = 10), Two-sample T test, t = 3.624, df = 18, 95% CI: 10.93 to 41.07, **$P = 0.0019$, compared with control mice. **I.** The time of latency to fall in the rotarod test (Con: N = 10; CRS: N = 10), Two-sample T test, t = 0.2178, df = 18, 95% CI:-11.24 to 13.84, $P = 0.8300$, compared with control mice. **J.** Representative Western blot assay (i) and summarized data (ii) showing the expression of TRPC6 and GAPDH in the VTA of the CRS mice (Con: N = 10; CRS: N = 14). Two-sample T test, t = 7.369, df = 22, 95% CI: -0.8006 to -0.4489, $****P < 0.0001$. $**P < 0.01$, $***P < 0.001$, $****P < 0.0001$. N is the number of mice used.

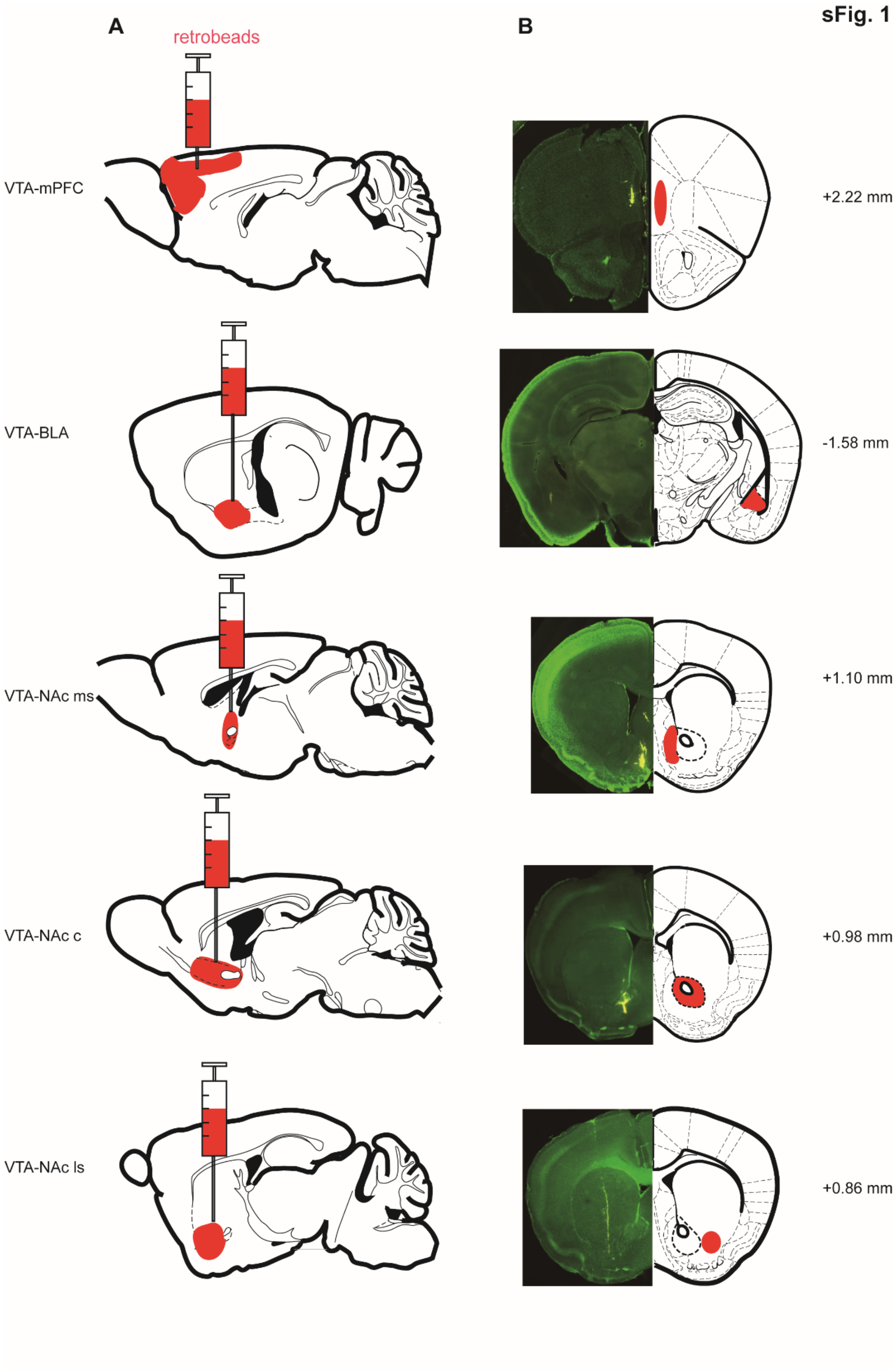
sFig. 1
A
retrobeads
B
VTA-mPFC
+2.22 mm
VTA-BLA
-1.58 mm
VTA-NAc ms
+1.10 mm
VTA-NAc c
+0.98 mm
VTA-NAc ls
+0.86 mm

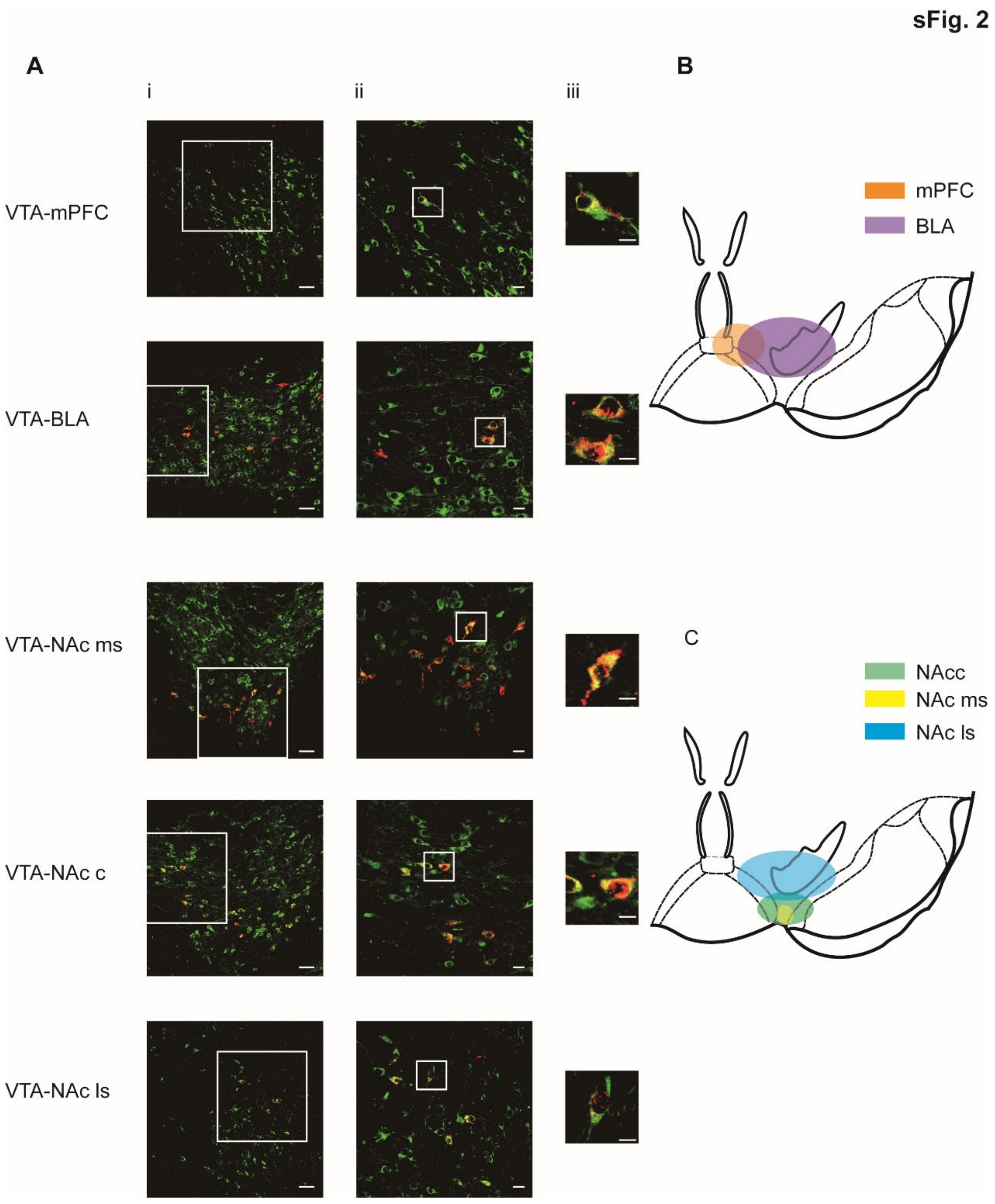

sFig. 2
A
i
ii
iii
VTA-mPFC
VTA-BLA
VTA-NAc ms
VTA-NAc c
VTA-NAc ls
B
mPFC
BLA
C
NAcc
NAc ms
NAc ls


169

sFig. 3

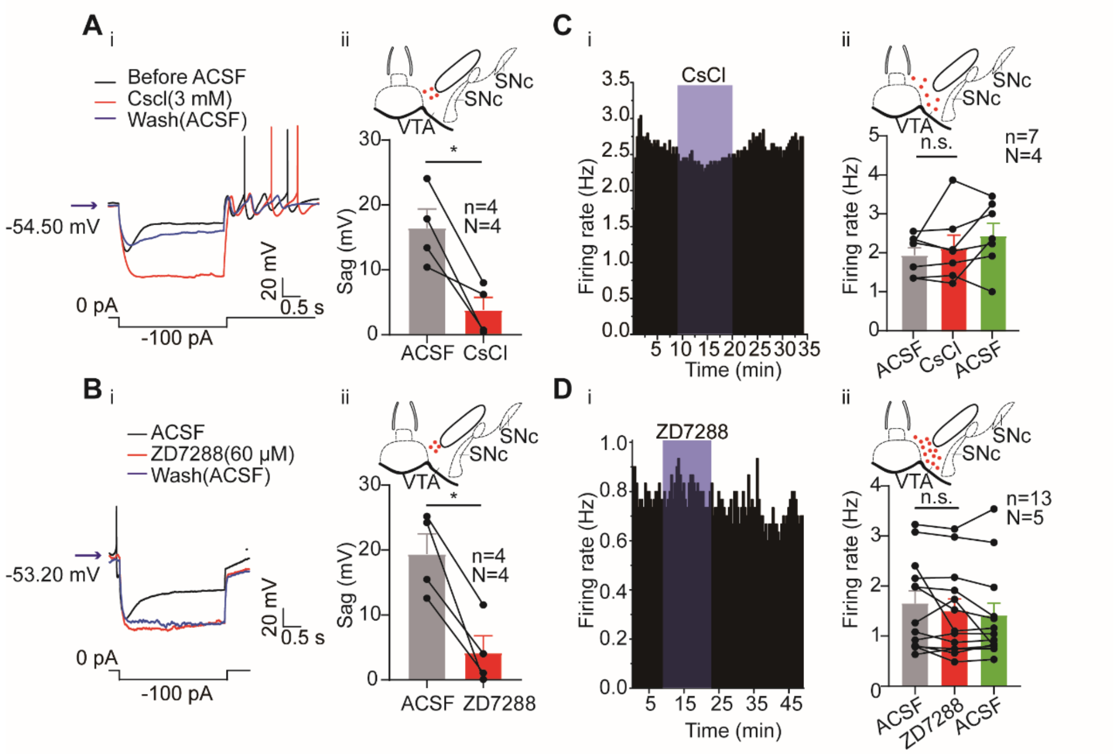
A i
Before ACSF
Cscl(3 mM)
Wash(ACSF)
-54.50 mV
20 mV
0.5 s
0 pA
-100 pA
ii
SNc
SNc
VTA
Sag (mV)
n=4
N=4
ACSF
CsCl
C i
CsCl
Firing rate (Hz)
Time (min)
ii
n.s.
n=7
N=4
Firing rate (Hz)
ACSF
CsCl
ACSF
B i
ACSF
ZD7288(60 μM)
Wash(ACSF)
-53.20 mV
20 mV
0.5 s
0 pA
-100 pA
ii
Sag (mV)
n=4
N=4
ACSF
ZD7288
D i
ZD7288
Firing rate (Hz)
Time (min)
ii
n.s.
n=13
N=5
Firing rate (Hz)
ACSF
ZD7288
ACSF

170

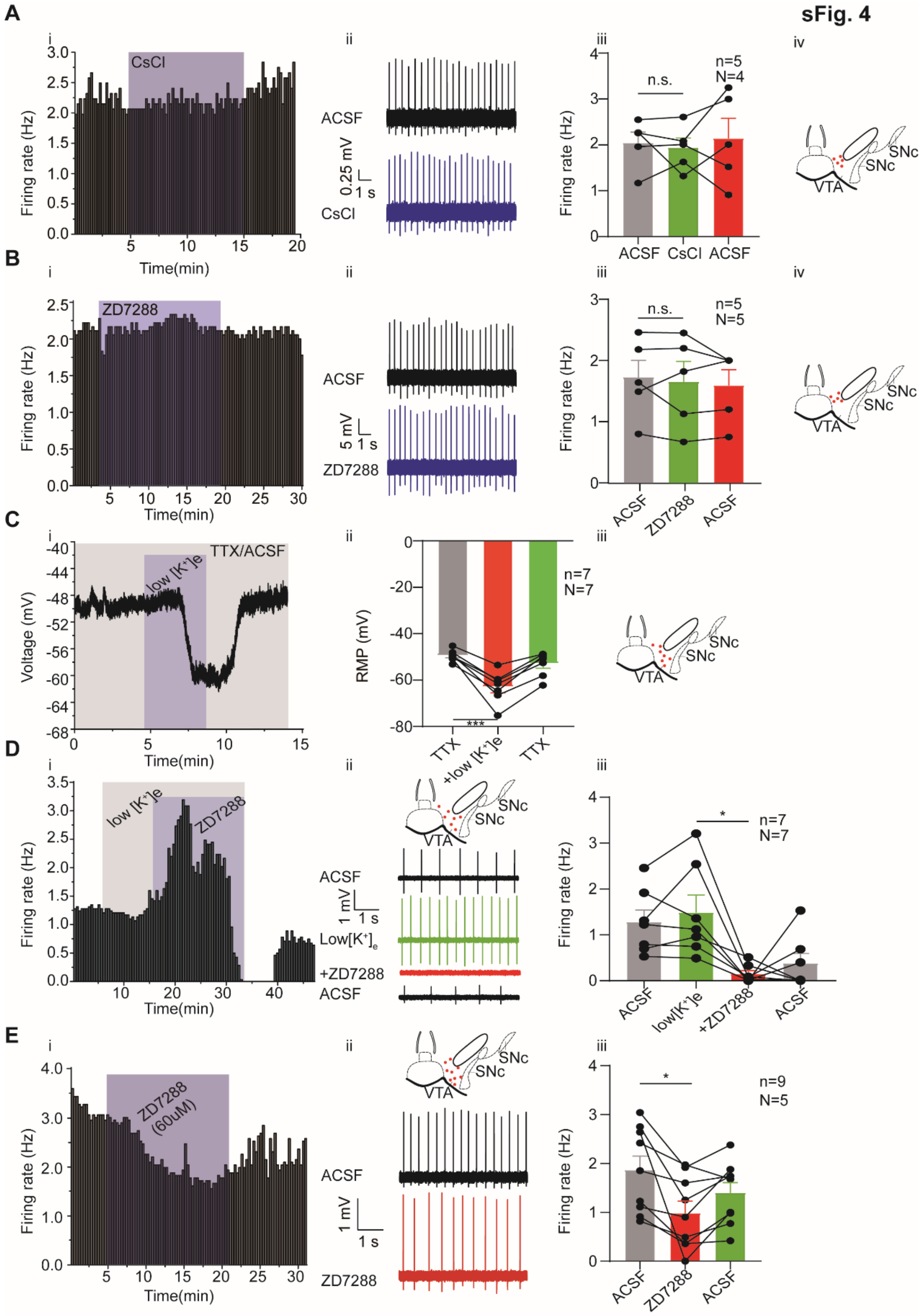

sFig. 4
A
CsCl
Firing rate (Hz)
Time(min)
ACSF
CsCl
0.25 mV
1 s
n.s.
n=5
N=4
ACSF CsCl ACSF
SNc
SNc
VTA
B
ZD7288
ACSF
5 mV
1 s
n=5
N=5
C
TTX/ACSF
low [K+]e
Voltage (mV)
RMP (mV)
n=7
N=7
***
TTX
+low [K+]e
D
low [K+]e
ZD7288
Low[K+]e
+ZD7288
*
E
ZD7288 (60uM)
1 mV
n=9
N=5

sFig. 5

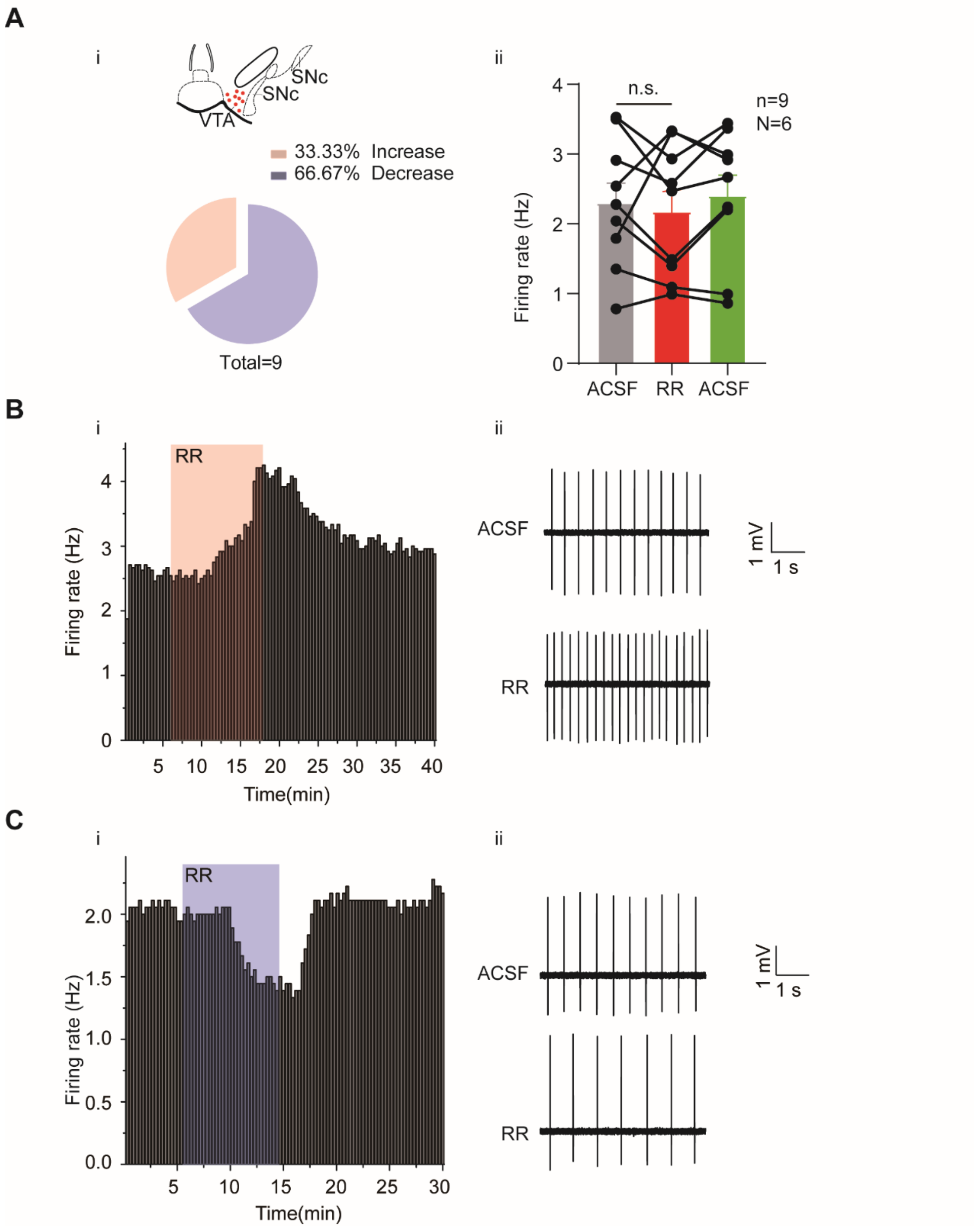
A
i
SNc
SNc
VTA
33.33% Increase
66.67% Decrease
Total=9
ii
n.s.
n=9
N=6
Firing rate (Hz)
ACSF
RR
ACSF
B
i
RR
Firing rate (Hz)
Time(min)
ii
ACSF
RR
1 mV
1 s
C
i
RR
Firing rate (Hz)
Time(min)
ii
ACSF
RR
1 mV
1 s

**sFig. 6**

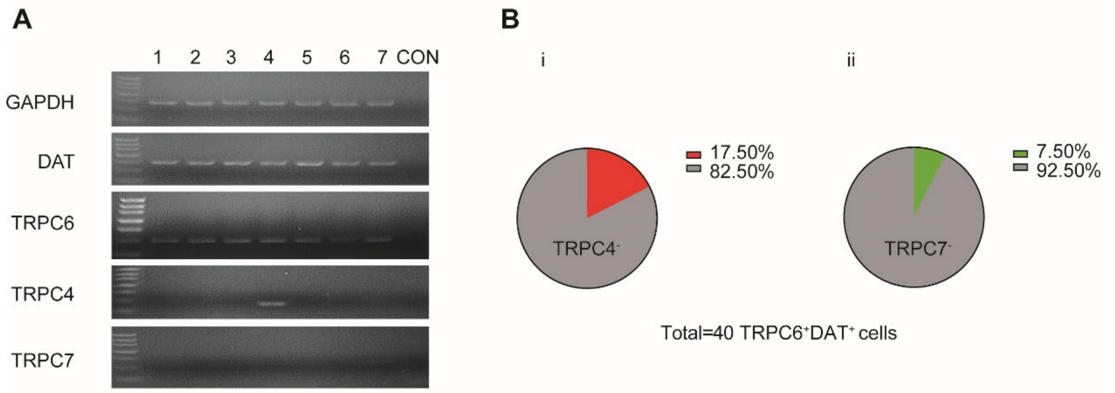
A
1 2 3 4 5 6 7 CON
GAPDH
DAT
TRPC6
TRPC4
TRPC7
B
i
ii
17.50%
82.50%
TRPC4-
7.50%
92.50%
TRPC7-
Total=40 TRPC6+DAT+ cells

**sFig. 7**

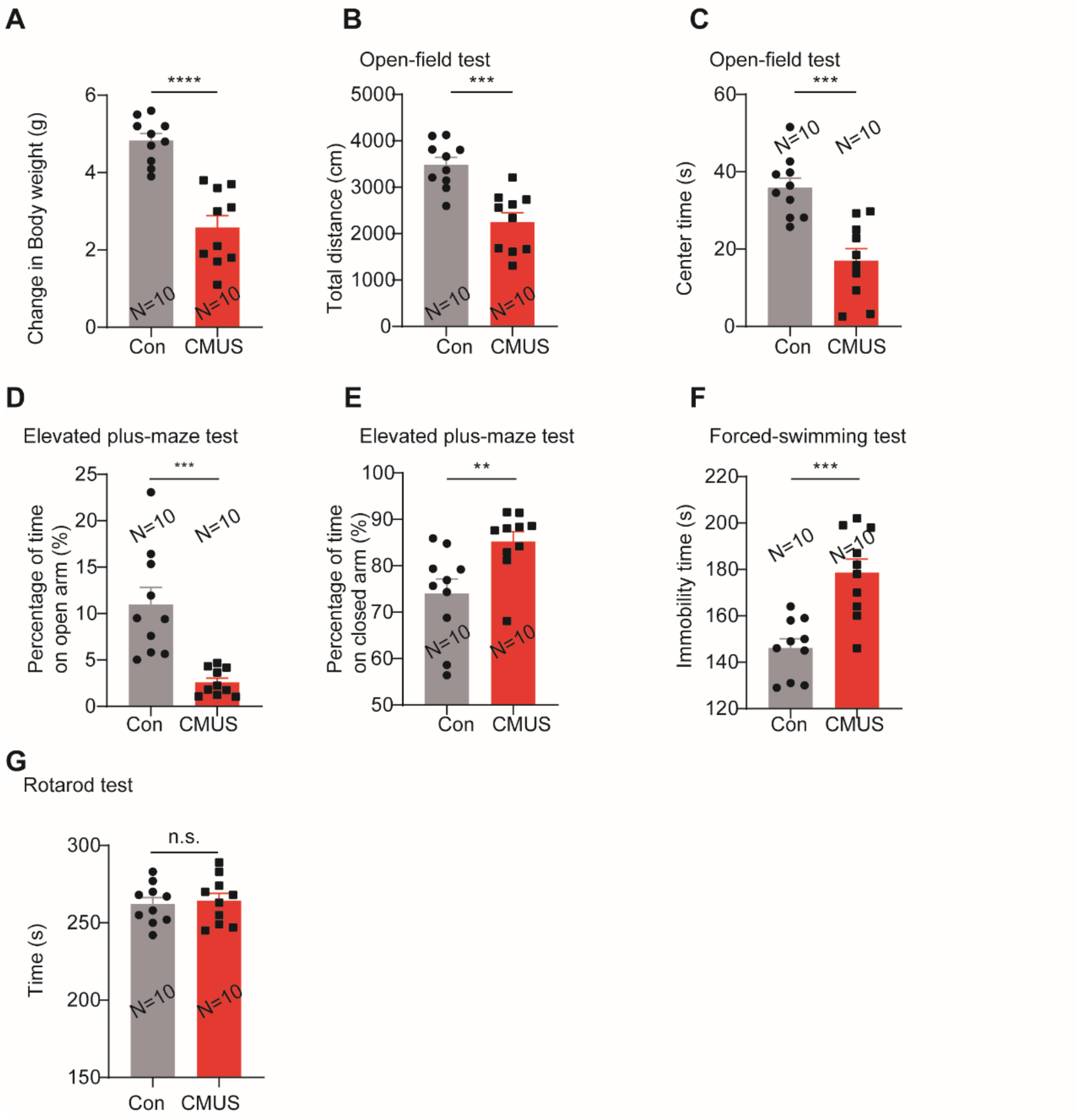
A
Change in Body weight (g)
****
N=10
N=10
Con
CMUS
B
Open-field test
Total distance (cm)
***
N=10
N=10
Con
CMUS
C
Open-field test
Center time (s)
***
N=10
N=10
Con
CMUS
D
Elevated plus-maze test
Percentage of time on open arm (%)
***
N=10
N=10
Con
CMUS
E
Elevated plus-maze test
Percentage of time on closed arm (%)
**
N=10
N=10
Con
CMUS
F
Forced-swimming test
Immobility time (s)
***
N=10
N=10
Con
CMUS
G
Rotarod test
Time (s)
n.s.
N=10
N=10
Con
CMUS

sFig. 8

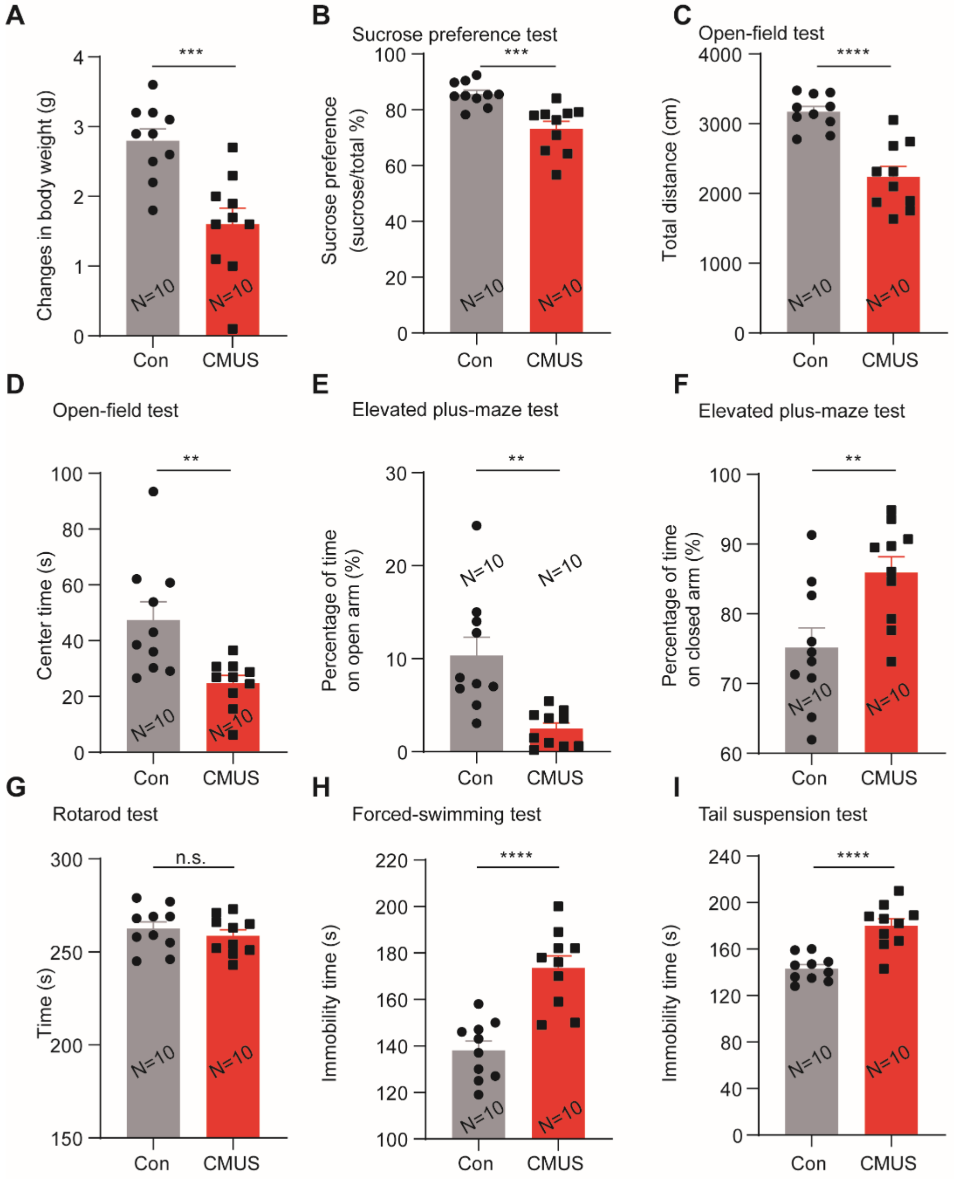


175

sFig. 9

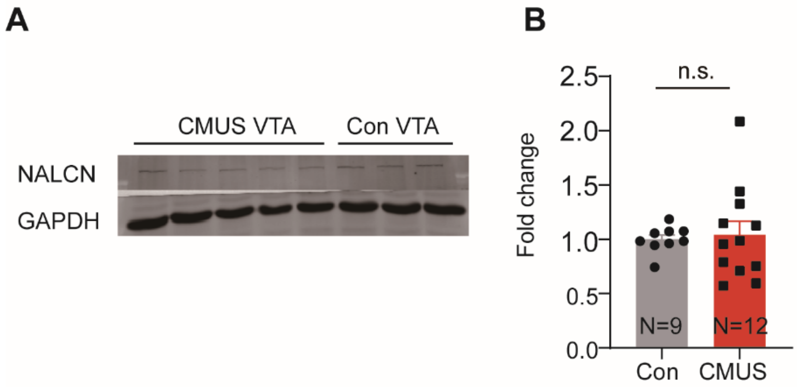


176

**sFig. 10**

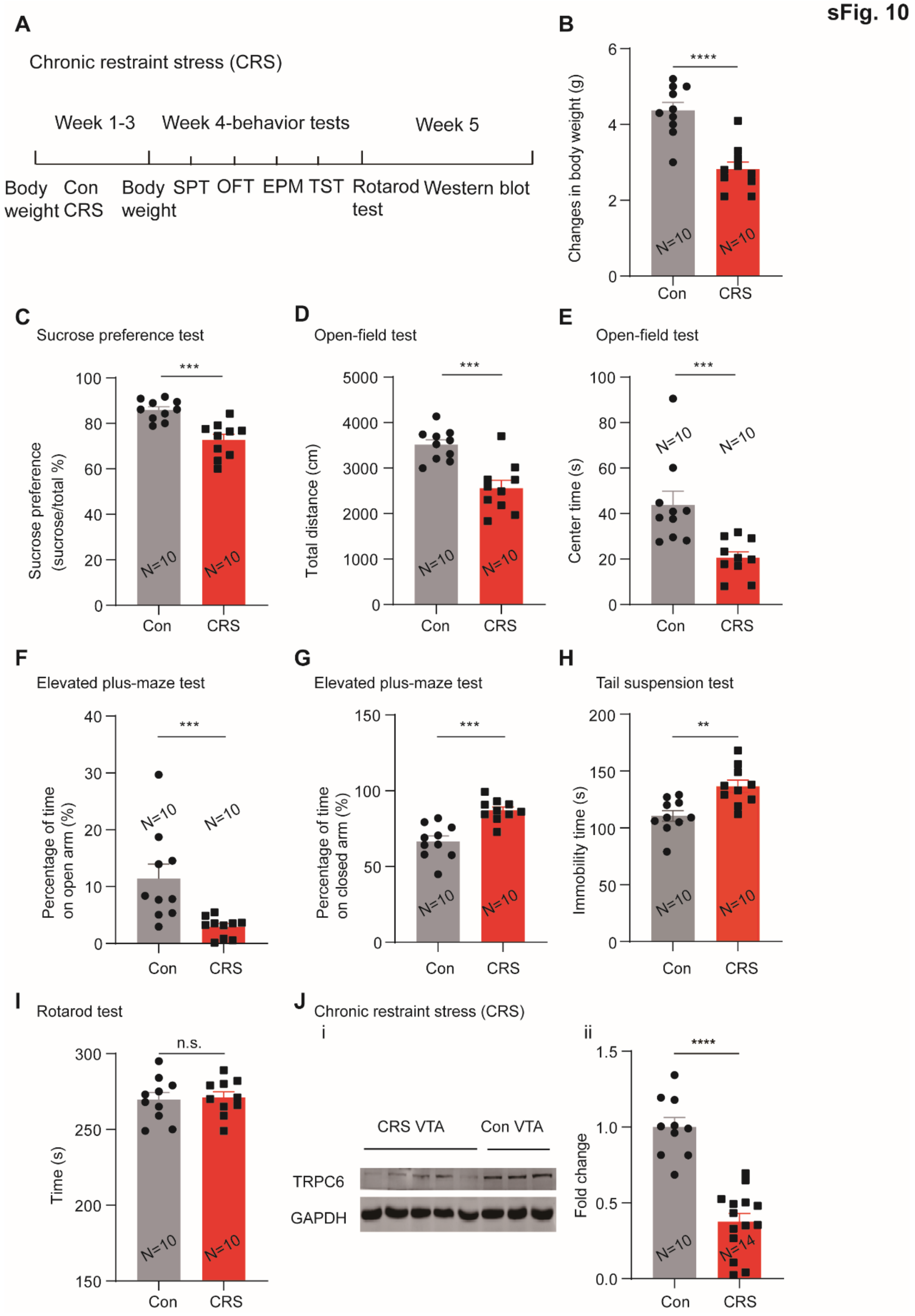
A
Chronic restraint stress (CRS)
Week 1-3
Week 4-behavior tests
Week 5
Body weight
Con CRS
Body weight
SPT
OFT
EPM
TST
Rotarod test
Western blot
B
Changes in body weight (g)
****
N=10
N=10
Con
CRS
C
Sucrose preference test
Sucrose preference (sucrose/total %)
***
D
Open-field test
Total distance (cm)
***
E
Open-field test
Center time (s)
***
F
Elevated plus-maze test
Percentage of time on open arm (%)
***
G
Elevated plus-maze test
Percentage of time on closed arm (%)
***
H
Tail suspension test
Immobility time (s)
**
I
Rotarod test
Time (s)
n.s.
J
Chronic restraint stress (CRS)
i
CRS VTA
Con VTA
TRPC6
GAPDH
ii
Fold change
****
N=14
